# A T7 phage factor required for managing RpoS in *Escherichia coli*

**DOI:** 10.1101/262279

**Authors:** Aline Tabib-Salazar, Bing Liu, Declan Barker, Lynn Burchell, Udi Qimron, Steve J. Matthews, Sivaramesh Wigneshweraraj

**Affiliations:** BioBank, First Affiliated Hospital, School of medicine, Xi’an Jiaotong University, Shaanxi Sheng, China; MRC Centre for Molecular Bacteriology and Infection, Imperial College London, London, SW7 2AZ, UK; Department of Clinical Microbiology and Immunology, Sackler School of Medicine, Tel Aviv University, Tel Aviv 69978, Israel.

**Keywords:** *Escherichia coli*, RNA polymerase, T7 phage, Transcription regulation, Stationary phase

## Abstract

T7 development in *Escherichia coli* requires the inhibition of the housekeeping form of the bacterial RNA polymerase (RNAP), Eσ^70^, by two T7 proteins: Gp2 and Gp5.7. While the biological role of Gp2 is well understood, that of Gp5.7 remains to be fully deciphered. Here, we present results from functional and structural analyses to reveal that Gp5.7 primarily serves to inhibit Eσ^S^, the predominant form of the RNAP in the stationary phase of growth, which accumulates in exponentially growing *E. coli* as a consequence of buildup of guanosine pentaphosphate ((p)ppGpp) during T7 development. We further demonstrate a requirement of Gp5.7 for T7 development in *E. coli* cells in the stationary phase of growth. Our finding represents a paradigm for how some lytic phages have evolved distinct mechanisms to inhibit the bacterial transcription machinery to facilitate phage development in bacteria in the exponential and stationary phases of growth.

**Significance statement:** Virus that infect bacteria (phages) represent the most abundant living entities on the planet and many aspects of our fundamental knowledge of phage-bacteria relationships have been derived in the context of exponentially growing bacteria. In the case of the prototypical *Escherichia coli* phage T7, specific inhibition of the housekeeping form of the RNA polymerase (Eσ^70^) by a T7 protein, called Gp2, is essential for the development of viral progeny. We now reveal that T7 uses a second specific inhibitor that selectively inhibits the stationary phase RNAP (Eσ^S^), which enables T7 to develop well in exponentially growing and stationary phase bacteria. The results have broad implications for our understanding of phage-bacteria relationships and therapeutic application of phages.

---

Viruses of bacteria, phages, have evolved diverse and sophisticated mechanisms to takeover essential host processes to facilitate the successful development of phage progeny. Many such host takeover mechanisms involve small proteins that interact with and repurpose, inhibit or modulate the activity of essential bacterial enzymes, which as a consequence, often result in the demise of the bacterial cell (1). Thus, a detailed understanding of phage-encoded antibacterial small proteins and their bacterial targets at a molecular level will not only unravel new phage biology, but may also inform and inspire the discovery of novel antibacterial targets and antibacterial compounds. Unsurprisingly, the acquisition of the bacterial transcription machinery, the RNA polymerase (RNAP), is a major mechanism by which phages reprogram bacterial cellular processes in order to mount a successful infection (2, 3). The prototypical lytic phage of *Escherichia coli*, T7, synthesizes three proteins, Gp0.7, Gp2 and Gp5.7, that interact with host RNAP, to facilitate the temporal coordinated expression of its genome. The genes of T7 are categorized as early, middle and late to reflect the timing of their expression during the infection process. Early and middle genes generally encode proteins required for phage RNA synthesis, DNA replication and host takeover, whereas the late genes specify T7 virion assembly and structural proteins. The translocation of the T7 genome into *E. coli* is a transcription-coupled process and requires the housekeeping form of the host RNAP (Eσ^70^) to transcribe the early genes from three strong early gene promoters, T7 A1, A2 and A3, and catalyze the entry of T7 DNA into the cell (4). The coordinated action of the early gene product Gp0.7 and the essential middle gene product Gp2 subsequently shuts off Eσ^70^ activity on the T7 genome. The viral single-subunit RNAP (T7 RNAP, Gp1, a product of an early gene) transcribes the middle and late viral genes. The shutting down of host RNAP is crucial for the coordination of the activities of bacterial and phage RNAPs on the phage genome and thus consequently for successful completion of the infection cycle: Gp0.7 is a protein kinase that phosphorylates Eσ^70^, leading to increased transcription termination at sites located between the early and middle genes on the T7 genome (5, 6); Gp2 binds in the main DNA binding channel of Eσ^70^ and thereby prevents the formation of the transcriptionally-proficient open promoter complex (RP_O_) at the T7 A1-3 promoters (7). Gp2 is indispensable for T7 growth. In a T7 Δ*gp2* phage, aberrant transcription of middle and late T7 genes (that are normally transcribed by the T7 RNAP) by Eσ^70^ results in interference between the two RNAPs and, consequently, in aborted infection (5). Recently, a T7 middle gene product, Gp5.7, was identified as a repressor of RP_O_ formation specifically on the T7 A1-3 promoters by Eσ^70^ molecules, which might have escaped inhibition by Gp2 (8). However, since phage genomes tend to be compact and efficient, it is puzzling that T7 has evolved two markedly different proteins to inhibit Eσ^70^, especially since Gp5.7, unlike Gp2, is a relatively poor inhibitor of Eσ^70^ (8). In this study, we unveil additional biological roles for Gp5.7 during T7 development in *E. coli*.

## Results

### Gp5.7 is an inhibitor of the *E. coli* stationary phase RNAP, Eσ^S^

Previously, we posited that Gp5.7 prevents transcription initiation from T7 A1-A3 promoters by Eσ^70^ that might have escaped inhibition by Gp2 (8). While this still remains a role for Gp5.7 in T7 development in *E. coli*, we noted a report by Friesen and Fill describing the accumulation of the stress signaling nucleotide guanosine pentaphosphate, (p)ppGpp, in a valine auxotroph strain of *E. coli* during T7 development (9). Since (p)ppGpp simultaneously induces σ^S^ transcription and accumulation of σ^S^ (the predominant σ factor active in stationary phase *E. coli*) and, considering that Gp5.7 is an inefficient inhibitor of Eσ^70^ compared to Gp2 (8), we contemplated whether Gp5.7 might preferentially inhibit Eσ^S^ over Eσ^70^. Initially, we established that (p)ppGpp does indeed accumulate during T7 development in exponentially growing wild-type *E. coli* cells (Fig. 1A). We further demonstrated that the accumulation of (p)ppGpp is accompanied by an increase in the intracellular levels of σ^S^ during T7 development in exponentially growing *E. coli* (Fig. 1B). Control experiments with a *relA*/*spoT* mutant *E. coli* strain confirmed that the accumulation of σ^S^ during T7 infection was indeed (p)ppGpp-dependent (Fig. 1B).

**Fig. 1.**
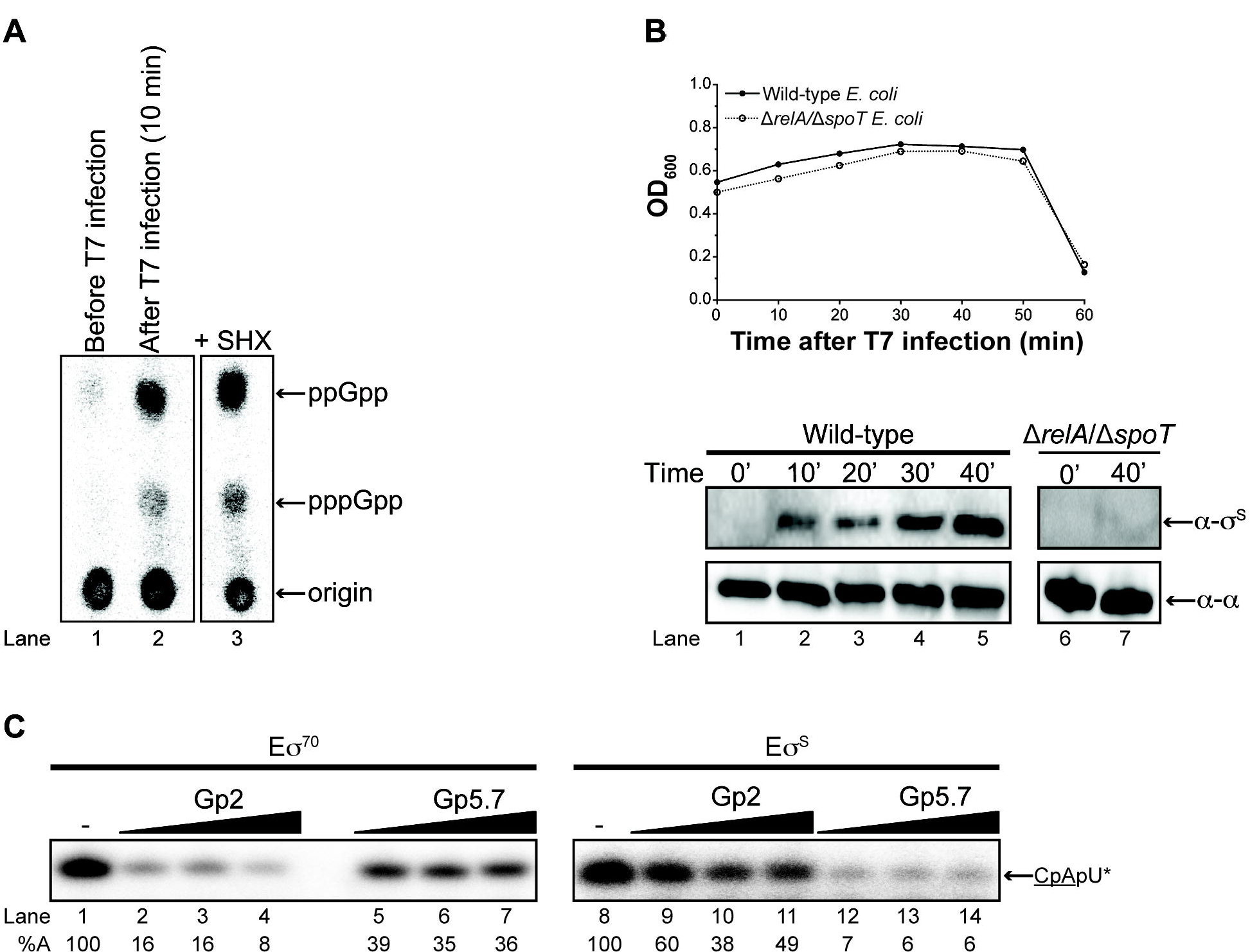
Gp5.7 is an inhibitor of the *E. coli* stationary phase RNAP, Eσ^S^. **(A)** PEI cellulose showing (p)ppGpp production during T7 infection. Lane 1: before T7 infection, lane 2: 10 min after infection with T7 and lane 3: positive control showing (p)ppGpp production in response to addition of sodium hydroxamate (SHX). The migration positions of ppGpp and (p)ppGpp and the origin where the samples were spotted are indicated. **(B)** Expression of σ^S^ during T7 infection. *Top*: Graph showing the optical density (OD_600nm_) of wild-type and Δ*relA*/Δ*spoT E. coli* cultures as a function of time after infection with T7 phage. *Bottom*: Image of a Western blot probed with anti-σ^S^ and anti-RNAP α-subunit (loading control) antibodies. Lanes 1 to 5 contain whole cell extracts of wild-type *E. coli* cells at 0, 10, 20, 30 and 40 min after infection with T7; lanes 6 and 7 contain whole cell extracts of Δ*relA*/Δ*spoT E. coli* cells at 0 and 40 min after infection with T7. **(C)** Autoradiograph of denaturing gels comparing the ability of Eσ^S^ and Eσ^70^ to synthesize a dinucleotide-primed RNA product from the T7 A1 promoter in the absence and presence of Gp2 and Gp5.7. The dinucleotide used in the assay is underlined and the asterisks indicate the radiolabelled nucleotide. The concentration of Eσ^S^ and Eσ^70^ was 75 nM and Gp2 and Gp5.7 were present at 75, 150 and 300 and 1200, 1500 and 1875 nM, respectively. The percentage of RNA transcript synthesized (%A) in the reactions containing Gp5.7 or Gp2 with respect to reactions with no Gp5.7 or Gp2 added is given at the bottom of the gel and the value obtained in at least three independent experiments fell within 3– 5% of the %A value shown.

Next, we tested whether Eσ^S^ could initiate transcription form the T7 A1 promoter as efficiently as Eσ^70^. To do this, we conducted an *in vitro* transcription assay using a 65-bp DNA fragment containing the T7 A1 promoter sequence as the template. Under the conditions used here, this assay reports the ability of Eσ^70^ or Eσ^S^ to bind to the promoter, initiate DNA strand separation, and synthesize a trinucleotide RNA transcript, CpApU, which is complementary to the first three nucleotides (+1 to +3) of the sequence of the template strand of the T7 A1 promoter. The results shown in Fig. 1C revealed that Eσ^S^ could initiate transcription form the T7 A1 promoter as efficiently as Eσ^70^ (Fig. 1C, compare lanes 1 and 8). Consistent with previous results (7, 10), Gp2 inhibited Eσ^70^ activity by >80% when present at a molar ratio of 1:1 with respect to Eσ^70^; in contrast, Gp2 inhibited Eσ^S^ by only 40% when present at a molar ratio of 1:1 with respect to Eσ^S^ (Fig. 1C, compare lanes 2 and 9). However, as previously shown (8), ~16-fold more Gp5.7 than Gp2 was required to obtain a ~60% inhibition of Eσ^70^ (Fig. 1C, compare lanes 2 and 5). Strikingly, with the same concentration of Gp5.7, we observed >90% inhibition of Eσ^S^ activity (Fig. 1C, lane 12). Control experiments with a functionally defective mutant of Gp5.7 (Gp5.7-L42A (8)) confirmed that the inhibition of Eσ^S^ activity on the T7 A1 promoter by Gp5.7 was specific (Fig. S1). It thus seems that Gp5.7 is a more efficient inhibitor of Eσ^S^ than of Eσ^70^ (Fig. 1C). In contrast and consistent with previous observations (7), Gp2 is a more effective inhibitor of Eσ^70^ than Eσ^S^ (Fig. 1C). Overall, we conclude that Gp5.7 and Gp2 are both required to fully shutdown the Eσ^70^ and Eσ^S^ to allow optimal T7 development in *E. coli* cells during the exponential phase of growth.

### Gp5.7 interacts with region 4 of σ^S^

Although Gp5.7 interacts with the core subunits of the RNAP (8), our results indicate that σ^S^ would likely constitute a major interacting site of Gp5.7. Therefore, we next focused on identifying the Gp5.7 interacting site on σ^S^. Nickel affinity pull-down experiments with hexa-histidine tagged σ^S^ (6xHis-σ^S^) and untagged Gp5.7 were conducted. As shown in Fig. 2A, incubation of 6xHis-σ^S^ (lane 2) with whole *E. coli* cell extracts in which untagged Gp5.7 is overexpressed from a plasmid (lane 3) led to the co-purification of untagged Gp5.7 with 6xHis-σ^S^ (lane 5). Control experiments with *E. coli* whole cell extracts with an empty plasmid (Fig. 2A, lanes 4 and 6) confirmed that Gp5.7 interacts specifically with 6xHis-σ^S^ and co-purifies with it. Since we showed previously that Gp5.7 interacts with promoter DNA proximal to or overlapping the consensus −35 motif of the T7 A1 promoter (8), which is also bound by the conserved region 4 (R4) domain of σ factors, we considered whether R4 domain of σ^S^ could be a binding site for Gp5.7. To test this, we conducted affinity pull-down experiments as in Fig. 2A using a hexa-histidine tagged version of the R4 domain (amino acid residues 245-330) of σ^S^ (6xHis-σ^S^R4). As shown in Fig. 2B (lane 5), we detected untagged Gp5.7 co-purifying with the 6xHis-σ^S^R4 domain. In the converse experiment, we repeated affinity pull-down experiments as in Fig. 2A using FLAG epitope tagged σ^S^ lacking the R4 domain (FLAG-σ^S^ΔR4). As indicated in Fig. 2C, and as expected, we failed to detect untagged Gp5.7 co-purifying with the FLAG-σ^S^ΔR4 protein (lane 5). Based on the affinity pull-down experiments shown in Fig. 2A-C, we conclude that the R4 domain of σ^S^ constitutes the binding site for Gp5.7.

**Fig. 2.**
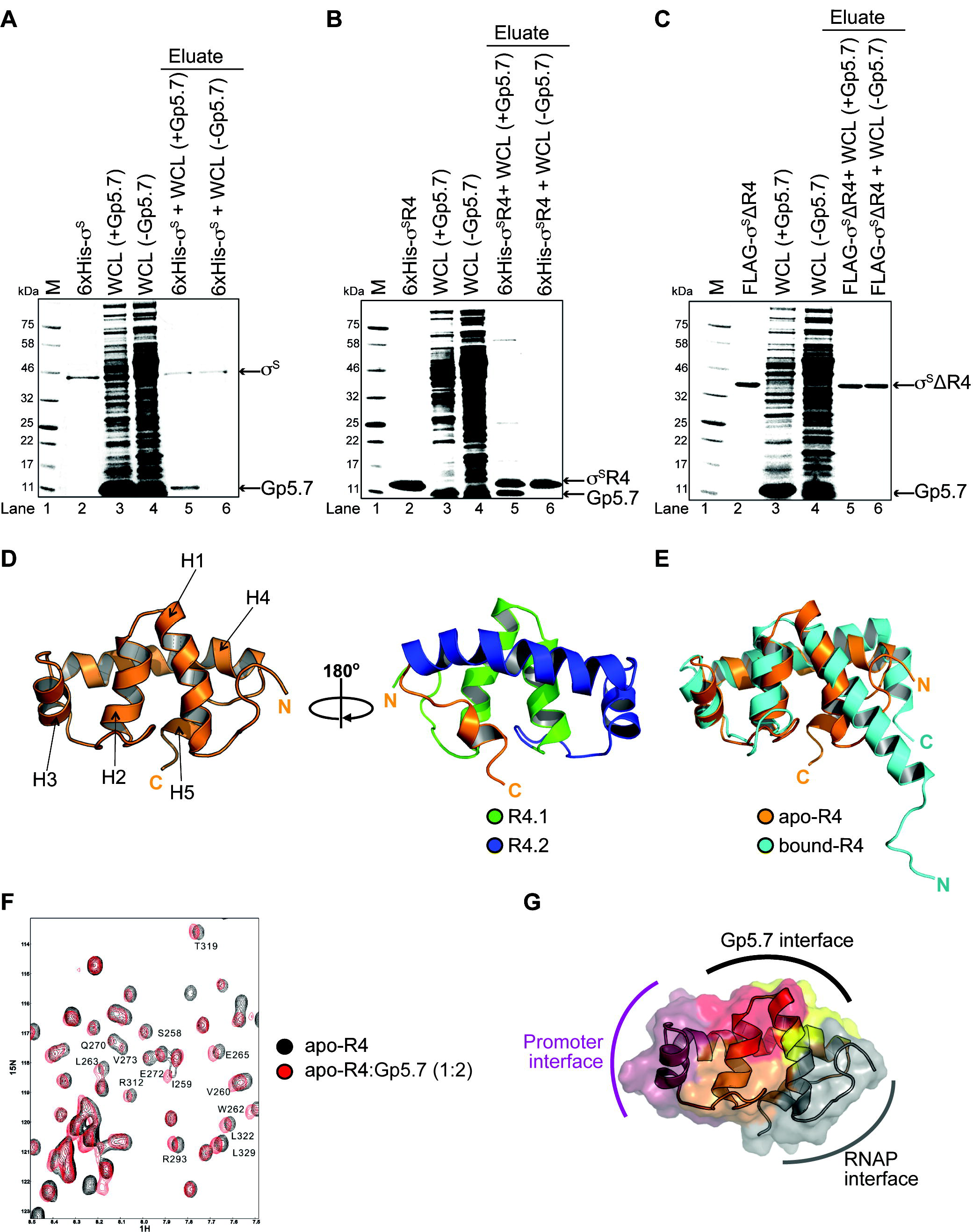
Structural insights into the interaction between R4 domain of σ^S^ and Gp5.7. **(A)** Image of a denaturing gel showing that 6xHis-σ^S^ pulls down overexpressed untagged Gp5.7 from *E. coli* whole cell lysate. The migration positions of the Gp5.7 and 6xHis-σ^S^ are indicated. **(B)** As in (A), but using 6xHis-σ^S^R4 domain. **(C)** As in (A), but using FLAG-σ^S^ΔR4. **(D)** *Left*: Cartoon representation of apo-σ^S^R4 solution structure. *Right:* The image on the left rotated 180° clockwise with the α-helices corresponding to R4 sub-regions 4.1 (in green) and 4.2 (in blue) indicated. **(E)** Overlay of cartoon representations of apo-σ^S^R4 domain (orange) and bound-σ^S^R4 domain (cyan). **(F)** Overlay of 2D ^1^H-^15^N HSQC spectra of the apo-σ^S^R4 without (black) and with Gp5.7 (red) recorded at pH 6.0, 300 K. Peaks that experienced broadening or chemical shift perturbation are labelled according to the amino acid residues in σ^S^. **(G)** A surface representation of apo-σ^S^R4 showing the regions interfacing with Gp5.7 (amino acid residues with significant peak broadening and chemical shift perturbation are shown in red, while those that experience moderate peak broadening and chemical shift perturbation are shown in orange and those that experience weak peak broadening and chemical shift perturbation are shown in yellow); amino acid residues associated with interacting with or proximal to the promoter DNA (raspberry) and RNAP subunits (grey) are also shown.

### Structural insights into the interaction between R4 domain of σ^S^ and Gp5.7

To independently verify that R4 domain of σ^S^ constitutes the binding site for Gp5.7 within Eσ^S^ and to map the Gp5.7 interface within R4 of σ^S^, we solved the solution structure of the isolated 6xHis-σ^S^R4 domain by NMR spectroscopy (Fig. 2D and Table S1). The structure demonstrates that the R4 domain of σ^S^ (hereafter referred to as the apo-R4 domain) is able to fold as an isolated subdomain, consisting of five α helices (H1-H5). The α helices H1-H5 superpose well with the equivalent region from the crystal structure of Eσ^S^ transcription initiation complex, in which the R4 domain (hereafter referred to as the bound-R4 domain) is connected to RNAP subunits via a long and flexible linker (11). Interestingly, the apo-R4 domain exhibits some conformational differences in the carboxyl (C) and amino (N) termini compared to the bound-R4 domain (Fig. 2E). The N-terminus of the apo-R4 domain, which in the bound-R4 domain is connected to Eσ^S^ via a flexible linker, appears more disordered. The C-terminus of the apo-R4 domain contacts the core of the structure making the apo-R4 domain more compact and stable in the absence of the remaining σ^S^ domains, RNAP subunits or promoter DNA. We next recorded the 2D ^1^H-^15^N HSQC NMR spectra to monitor the backbone amide chemical shift and line-width perturbations for ^15^N-labelled apo-R4 domain in the presence of up to a 2-fold molar excess of unlabelled Gp5.7. As shown in Fig. 2F, several peaks exhibited measurable broadening effects and the extent of broadening correlating with the amount of Gp5.7 added (Fig. 2F). The Gp5.7 interaction surface was mapped on the structure of the apo-R4 domain (Fig. 2G), revealing that the main interacting residues localise to the C terminal part of H1 and N terminal part of H2. This analysis suggests that Gp5.7 binds between the RNAP facing surface and the promoter-facing surface of R4 of σ^S^ (notably H3 of 4.2 sub-region of R4; Fig. 2D). Overall, the results from the affinity pull-down and structural analyses unambiguously indicate that the R4 of σ^S^ constitutes the binding site for Gp5.7 on σ^S^.

### Gp5.7 inhibits RP_O_ formation by Eσ^S^ on the T7 A1 promoter

The location of the Gp5.7 binding surface on σ^S^ implies that Gp5.7 has evolved to target a σ^S^ domain potentially important for transcription initiation from the T7 A1 promoter. Previous reports from several groups have implied that the interaction between R4 of σ^70^ and the consensus −35 promoter region is required for the stabilization of early intermediate promoter complexes *en route* to the RP_O_ at the T7 A1 promoter (12-15). Therefore, we considered whether Gp5.7 inhibits Eσ^S^-dependent transcription from the T7 A1 promoter by antagonizing RP_O_ formation. In agreement with this view, whereas Eσ^S^ reconstituted with σ^S^ΔR4 was able to initiate transcription from a prototypical Eσ^70^-dependent promoter i.e. *lac*UV5 (albeit at a lower efficiency compared to reactions with wild-type Eσ^S^), we failed to detect any transcription by Eσ^S^ΔR4 from the T7 A1 promoter (Fig. 3A). We then conducted electrophoretic gel mobility shift assays at 4°C (to detect initial RNAP-promoter complex formation) and at 37°C (to detect RPo formation) with Eσ^S^ΔR4 and a γ-^32^P-labelled T7 A1 probe to determine whether Eσ^S^ΔR4 was able to interact with the T7 A1 promoter to form the initial promoter complex or whether the initial promoter complex formed by Eσ^S^ΔR4 on the T7 A1 promoter was unable to isomerize into the RPo, respectively. Results shown in Fig. 3B, demonstrated that although Eσ^S^ and Eσ^S^ΔR4 formed the initial promoter complex on the T7 A1 promoter equally well (lanes 2-3 and lanes 5-6), those formed by Eσ^S^ΔR4 seem unable to isomerize to form the transcriptionally proficient RPo (lanes 8-9 and lanes 11-12). Consistent with this conclusion, *in vitro* transcription assays with a T7 A1 promoter probe containing an heteroduplex segment between positions −12 and +2 (to mimic the RP_O_) revealed that Eσ^S^ΔR4 is able to synthesize the CpApU transcript (Fig. 3C, compare lanes 1 and 2 with lanes 3 and 4). In further support of the view that Gp5.7 inhibits RP_O_ formation at the T7 A1 promoter, Eσ^S^ was able to synthesize the CpApU transcript in the presence of Gp5.7 when the latter was added to a pre-formed RP_O_ (i.e. when Gp5.7 was added to the reaction following pre-incubation of Eσ^S^ and the homoduplex T7 A1 promoter at 37°C) (Fig. 3D). To better understand how Gp5.7 inhibits RP_O_ formation at the T7 A1 promoter, we used the 2D ^1^H-^15^N HSQC NMR data of the interaction between Gp5.7 and R4 of σ^S^ (Fig. 2F and 2G), the solution structure of Gp5.7 (8) and the X-ray crystal structures of Eσ^S^-transcription initiation complex (TIC) (in which the interaction between R4 of σ^S^ and the consensus −35 promoter region is absent; (11)) and the *E. coli* Eσ^70^ TIC (in which the interaction between the consensus −35 promoter region and the R4 of σ^70^ is present; (16)) to construct a model of Gp5.7 bound Eσ^S^ TIC using HADDOCK (17). This model, shown in Fig. S2, suggests that Gp5.7 binds to Eσ^S^ in such an orientation that the positively changed side chains of amino acid residues R24 and R47 face the DNA region immediately adjacent to the consensus −35 motif of the T7 A1 promoter. Since efficient RP_O_ formation at the T7 A1 promoter depends on the interaction between R4 of σ^70^ (12-15) and σ^S^ (Fig. 3A and 3B) and the consensus −35 promoter region, we envisage a scenario where the interaction of Gp5.7 with this region of the T7 A1 promoter antagonizes the interactions between R4 and the consensus −35 motif that are required for efficient RP_O_ formation at this promoter. This view is also supported by our previous observation that apo Gp5.7 interacts, albeit weakly, with the region immediate upstream of the −35 motif of the T7 A1 promoter (8) and an alanine substitution at R24 (but not R47) render Gp5.7 inactive *in vivo* (8). Further, the model suggests that Gp5.7 is also proximal to core RNAP subunits (notably the β subunit), consistent with the finding that Gp5.7 can interact with the RNAP in the absence of any σ factors (8). Overall, we conclude that R4 of σ^S^ is important for RP_O_ formation at the T7 A1 promoter and Gp5.7 inhibits RP_O_ formation by Eσ^S^ during T7 development by interfering with the R4 of σ^S^.

**Fig. 3.**
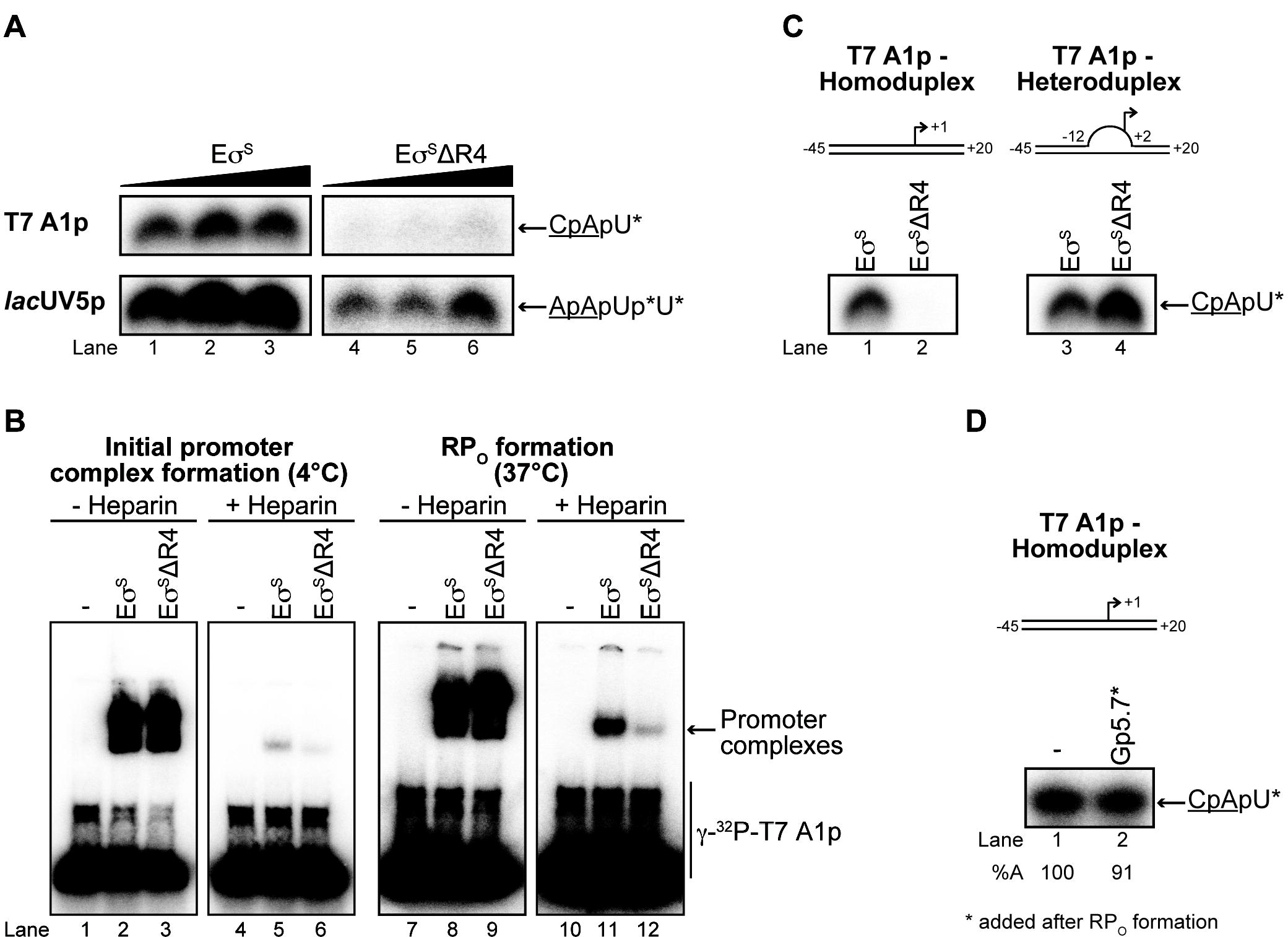
Gp5.7 inhibits RP_O_ formation by Eσ^S^ on the T7 A1 promoter. **(A)** Autoradiograph of denaturing gels showing the ability of the Eσ^S^ΔR4 to synthesize a dinucleotide-primed RNA product from the T7 A1 and *lac*UV5 promoters. The dinucleotide used in the assay is underlined and the asterisks indicate the radiolabeled nucleotide. **(B)** Autoradiographs of non-denaturing gels showing the ability of Eσ^S^ and Eσ^S^ΔR4 to form the initial promoter complex (< 4°C) and the RP_O_ (at 37°C) on the T7 A1 promoter. The migration positions of promoter complexes and free DNA are indicated. Reactions to which heparin were added are indicated. See methods for details. **(C)** As in (A) but using the T7 A1 homoduplex and heteroduplex (−12 to +2) promoters. **(D)** As in (A) but Gp5.7 (1875 nM) was added to the pre-formed RP_O_ (formed using 75 nM Eσ^S^ (see text for details)).

### A role for Gp5.7 in managing σ^S^ during T7 development in stationary phase *E. coli*

The results so far indicate that Eσ^S^ accumulates as a consequence of (p)ppGpp buildup during T7 development in exponentially growing *E. coli* cells and that Gp5.7 is required to preferentially inhibit Eσ^S^ activity on the T7 A1 promoter. However, when *E. coli* cells are in the stationary phase of growth, the major species of RNAP molecules will contain σ^S^ (also due to the buildup of (p)ppGpp in response to the stresses encountered by *E. coli* cells in the stationary phase of growth; reviewed in (18)). In addition, (p)ppGpp, together with Eσ^S^, also contributes to the shutting-down of cellular activities in the stationary phase of growth. Therefore, the development of a phage can be affected by changes in the growth state, and thus cellular activities, of the bacterial cell. Consistent with this view, Nowicki et al (19) recently reported that progeny production by Shiga toxin converting lamboid phages was significantly more efficient in a Δ*relA*/Δ*spoT E. coli* (which is unable to synthesize (p)ppGpp) than in its isogenic wild-type strain.

A phage plaque is a clearing in a bacterial lawn and plaques form via an outward diffusion of phage progeny virions that prey on surrounding bacteria. Therefore, the rate of plaque-enlargement can serve as a proxy for how efficiently a phage develops and produces progeny within an infected bacterial cell. Further, although the aging of the bacterial lawn often represents a major barrier for plaque-enlargement, T7 plaques have been reported to enlarge continually on matured *E. coli* lawns (20), suggesting that T7 has evolved specific mechanisms to infect and develop in the stationary phase of *E. coli* growth. Therefore, we investigated whether Gp5.7 is required for T7 development in *E. coli* in the stationary phase of growth by measuring plaque size formed on a lawn of *E. coli* as a function of incubation time in the context of wild-type and Δ*relA*/Δ*spoT E. coli* strains (recall accumulation of σ^S^ will be compromised in the mutant strain due to the absence of (p)ppGpp; see above (18)). As shown in Fig. 4, the rate of plaque-enlargement and plaque size on a lawn of wild-type *E. coli* infected with T7 wild-type and Δ*gp5.7* phage was indistinguishable for the first 12 hours (Fig. 4 and Movie S1). However, whereas T7 wild-type plaques continued to enlarge, the rate at which the plaques formed by T7 Δ*gp5.7* enlarged significantly slowed after ~12 hours of incubation and completely ceased after ~20 hours of incubation (Fig. 4 and Movie S1). Hence, after 72 hours of incubation, the size of the plaque formed by the T7 Δ*gp5.7* phage was ~50% smaller than the plaque formed by T7 wild-type phage on a lawn of wild-type *E. coli* cells. We were able to partially, yet specifically, revert the rate of plaque-enlargement and plaque size of the T7 Δ*gp5.7* phage to that of T7 wild-type phage by exogenously providing Gp5.7 from an inducible plasmid in *E. coli* (Fig. S3). The results overall imply that Gp5.7 is required for T7 development in *E. coli* in the stationary phase of growth. To independently verify this view, the relative efficiency of plaque formation (E.O.P) by the T7 Δ*gp5.7* phage on exponentially growing *E. coli* was compared with that on *E. coli* in the stationary phase of growth. Results shown in Fig. S4 indicated that the relative E.O.P by the T7 Δ*gp5.7* phage was almost 3 times lower than that of the wild-type phage (Fig. S4). This observation further underscores the view that T7 development in *E. coli* in the stationary phase of growth is compromised in the absence of Gp5.7.

**Fig. 4.**
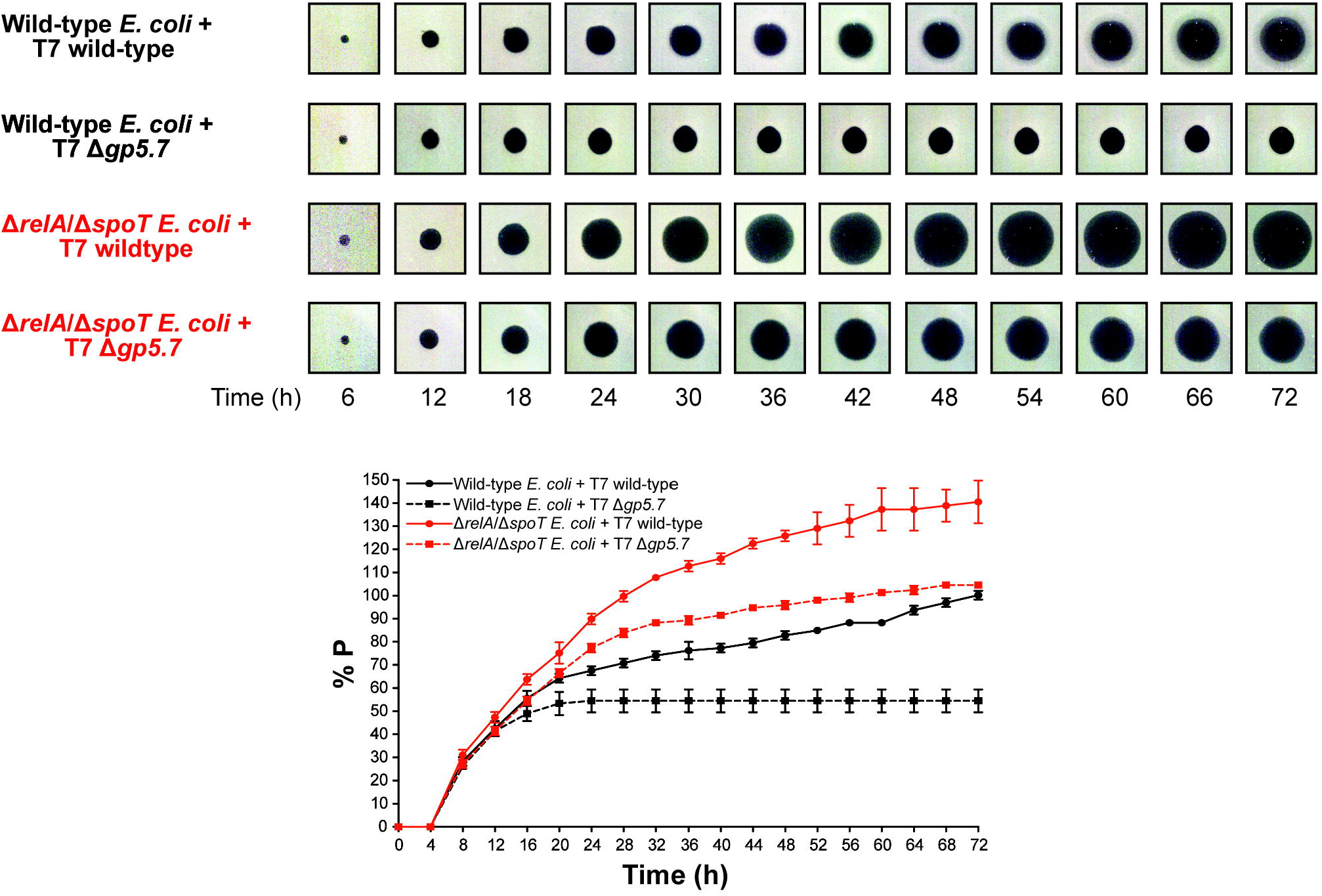
A role for Gp5.7 in managing σ^S^ during T7 development in stationary phase *E. coli*. *Top:* Representative scanned images of T7 wild-type and T7 Δ*gp5.7* plaques formed on a lawn of wild-type *E. coli* and Δ*relA*/Δ*spoT E. coli* over a 72 h incubation period. *Bottom:* Graph showing plaque size (%P) as percentage of final plaque size formed by T7 wild-type phage on a lawn of wild-type *E. coli* after 72 h of incubation (set at 100%) as a function of incubation time.

In marked contrast, although the rate of enlargement of the plaques formed by T7 wild-type phage was similar on a lawn of Δ*relA*/Δ*spoT E. coli* to that observed on a lawn of wild-type *E. coli* for the first 8 hours of incubation, the plaques formed by T7 wild-type phage continued to enlarge at a faster rate on a lawn of Δ*relA*/Δ*spoT E. coli* than on a lawn of wild-type *E. coli* lawn (Fig. 4). For example, after 48 hours of incubation, the size of the plaques formed by the T7 wild-type phage on a lawn of Δ*relA*/Δ*spoT E. coli* was ~2-fold larger than those formed on a lawn of wild-type *E. coli* (Fig. 4) Strikingly, whereas the plaques formed by the T7 Δ*gp5.7* phage ceased enlarging after ~20 hours of incubation on a lawn of wild-type *E. coli*, they continued to enlarge (albeit at a slower rate than that of T7 wild-type phage) on a lawn of Δ*relA*/Δ*spoT E. coli* (Fig. 4). Overall, although we cannot fully exclude possibility that the absence of (p)ppGpp in Δ*relA*/Δ*spoT E. coli* will generally provide a more favorable intracellular conditions for T7 development than in wild-type *E. coli* cells, the results clearly demonstrate that (i) the accumulation of (p)ppGpp during T7 infection antagonizes T7 development in *E. coli*; (ii) Gp5.7 represents a mechanism by which T7 overcomes the antagonistic effect of (p)ppGpp-mediated accumulation of σ^S^ on its development and therefore (iii) Gp5.7 is also a T7 factor required for T7 development in *E. coli* cells in the stationary phase of growth.

## Discussion

The inhibition of the host transcription machinery, the RNAP, is a central theme in the strategies used by phages to acquire their bacterial prey. In the prototypical *E. coli* phage T7, the switching from using the host RNAP for transcription of early T7 genes to the T7 RNAP for transcription of middle and late T7 genes is tightly regulated by two bacterial RNAP inhibitors: Gp2 and Gp5.7. Dysregulation of this process, for example due to the absence of any of these factors, is believed to lead to steric interference between the host and T7 RNAP molecules on the T7 genome and results in compromised or aborted development of phage progeny (5, 8). In an earlier study, we proposed that Gp5.7 acts as a ‘last line of defense’ molecule to prevent aberrant transcription of middle and late T7 genes by host RNAP molecules that may have escaped inhibition by Gp2 (8). The present study has uncovered additional biological roles for Gp5.7 in the T7 development cycle. The new biological roles for Gp5.7 in the T7 developmental cycle uncovered in the present study are summarized in the model in Fig. 5, which is partly supported by experimental evidence but also based on several assumptions (e.g. the differences in the intracellular levels of phage proteins and σ factors) that may not hold up as more evidence emerges. The results from the *in vitro* experiments presented here, infer that during infection of exponentially growing *E. coli* cells by T7 phage, Gp5.7 serves to inhibit transcription initiation from T7 A1-3 promoters by Eσ^S^ (Fig. 1C), which accumulates (Fig. 1B), possibly as a consequence of the (p)ppGpp-mediated stress response mechanism (Fig. 1A) to T7 infection (Fig. 5, box 1-3). Gp5.7 is also required for T7 development in *E. coli* in the stationary phase of growth (Fig. 5, box 4-6). In this case, we envisage that Gp5.7 will be absent when the transcription (of early T7 genes) dependent translocation of the T7 genome by Eσ^S^ occurs during infection of *E. coli* in the stationary phase of growth (Fig. 5, box 4), but becomes available when the Eσ^S^ is no longer required, i.e. when the T7 RNAP takes over the transcription of the middle and late genes (Fig. 5, box 5). The fact that Gp2 only poorly inhibits Eσ^S^ (7) further supports the need for Gp5.7 for T7 development in both exponentially growing and stationary phase *E. coli* cells. Thus, to the best of our knowledge, Gp5.7 is the only phage-encoded host RNAP inhibitor (or phage factor) described to date that is required for successful phage development in stationary phase bacteria. Intriguingly, we are unable to ‘rescue’ the T7 Δ*gp5.7* phage in a Δ*rpoS E. coli* strain in the context of the plaque-enlargement assay shown in Fig. 4. As shown in Fig. S5, wild-type and Δ*gp5.7* T7 phages are equally compromised to efficiently develop in the Δ*rpoS E. coli* strain. However, based on the assumption that the intracellular levels of Eσ^70^ will be higher in the Δ*rpoS E. coli* than in the wild-type *E. coli* due to the absence of the competing σ^S^ (21), we propose that T7 is unable to efficiently develop in the Δ*rpoS E. coli* because of inadequate ability of Gp2 and Gp5.7 (and Gp0.7) to inhibit the excess Eσ^70^ molecules (which will presumably dilute the intracellular pool of Gp2 and Gp5.7 (and Gp0.7) available to fully inhibit Eσ^70^ to allow optimal T7 development). Overall, our results indicate that, although T7 development in *E. coli* depends on the host RNAP (for transcription-dependent translocation of T7 genome into the bacterial cell and transcription of early T7 genes), efficient management of host RNAP activity is clearly obligatory for T7 to optimally develop both in exponentially growing and stationary phase *E. coli* cells. Consequently, any perturbations in RNAP levels or activity can have adverse effects on T7 development.

**Fig. 5.**
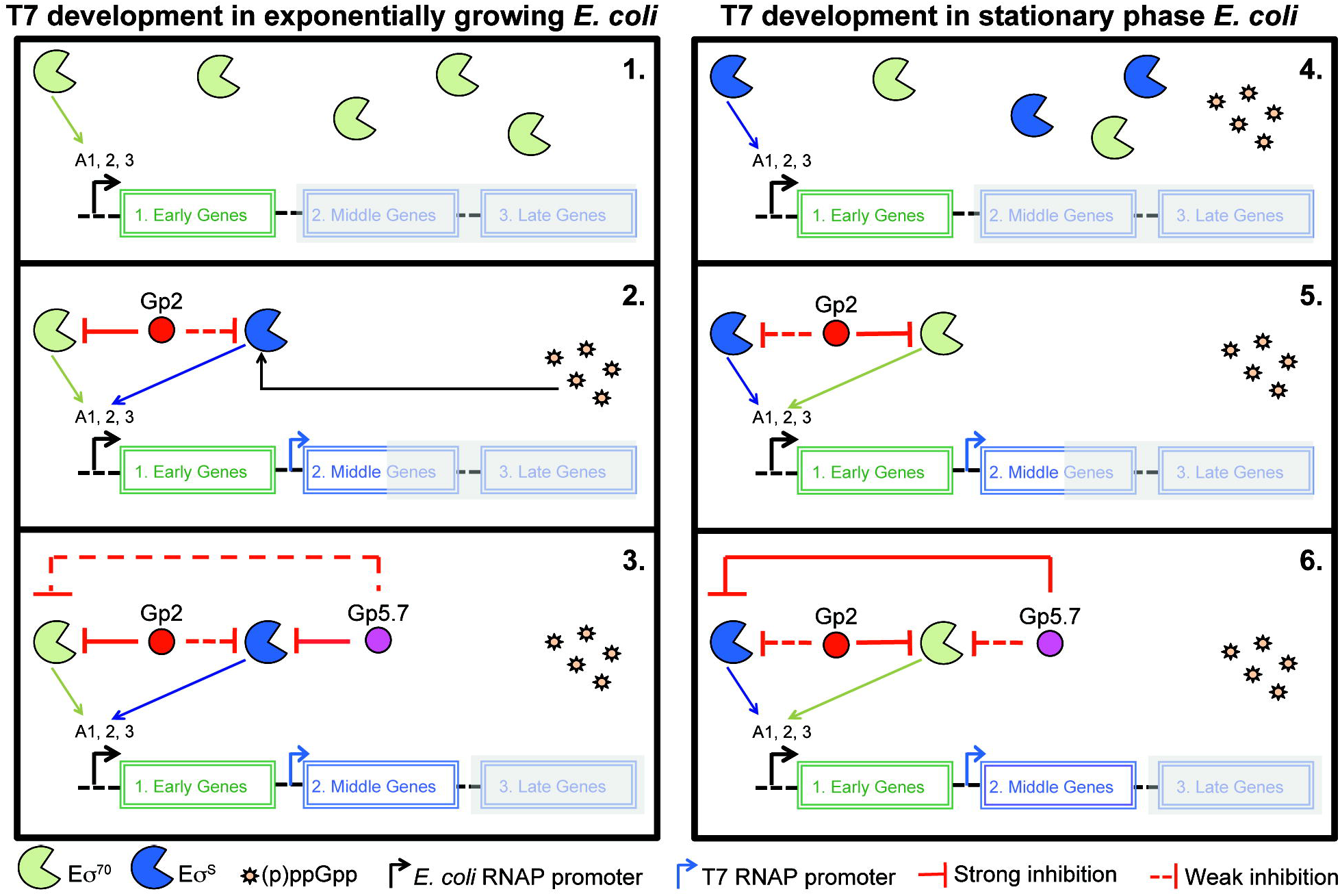
Model proposing how the host RNAP is managed during T7 development in exponentially growing and stationary phase *E. coli* cells.

We further note that, although Gp2 and Gp5.7 bind to sites located at different faces of the RNAP (with respect to the active center of the RNAP, the Gp2 binding site is located on the β’ jaw domain at the downstream face of the RNAP (10), whereas Gp5.7 binding site is located at the upstream face of the RNAP (this study)), both T7 proteins seem to inhibit RP_O_ formation by misappropriation of essential domains of the σ factor (region 1 of σ^70^ in case of Gp2 (7) (22) and R4 of σ^S^ in the case of Gp5.7 (this study)). Since the R4 domain of σ^70^ is also targeted by a T4 phage protein, called AsiA, to ‘recruit’ the host RNAP to transcribe phage genes (reviewed in (23)), it is interesting to speculate whether phages, regardless of their dependence on the host RNAP (unlike T7 phage, the T4 phage fully relies on the host RNAP for the transcription of its genes), have evolved to misappropriate essential bacterial σ factor domains to inhibit (e.g. T7) or redirect (e.g. T4) host RNAP activity to serve phage developmental requirements.

This study unambiguously shows that (p)ppGpp accumulates in T7 infected *E. coli* cells. The involvement of (p)ppGpp in phage development has been previously documented: For example, (p)ppGpp is required for the replication of phage Mu in *E. coli* ((24)) and in phage lambda it contributes to the switching between the lytic and lysogenic cycles (25). However, the role of (p)ppGpp in T7 development and the signaling pathway(s) that results in its synthesis are unknown. In *E. coli* two different pathways are involved in the production of (p)ppGpp: the RelA- and SpoT-dependent pathways (reviewed in (26)). RelA is associated with ribosomes and produces (p)ppGpp in response to uncharged tRNA in the ribosomal A-site during amino acid starvation. In contrast, SpoT is primarily responsible for the accumulation of (p)ppGpp in response to most stresses (e.g. fatty acid or iron starvation) and nutrient limitations (e.g. carbon starvation) apart from amino acid starvation. However, it seems paradoxical that T7 infected *E. coli* cells experience amino acid starvation since cellular translation becomes increased during T7 development (to serve phage gene expression needs) through the phosphorylation of translation elongation factors G, F and the ribosomal protein S6 (27). Although this study describes a strategy T7 uses to mitigate the effect of accumulation of (p)ppGpp during T7 development in *E. coli*, clearly, the role of (p)ppGpp in T7 development and the signaling pathway(s) that induce its synthesis warrant further investigation.

In summary, our study has uncovered a new aspect of T7 biology and the distinct strategies used by this phage to shut down bacterial RNAP for an optimal infection outcome in *E. coli* in exponentially growing and stationary phase of growth. The latter is clearly relevant to bacteria encountered by T7 in the natural environment, which are often in a starved and thus in a growth-attenuated or slow growing state. Therefore, the insights from this study also have implications on the emerging interest in the use of phages, phage-derived antibacterial compounds and their bacterial targets to treat bacterial infections where bacteria largely exist in a ‘stressed’ state and mostly depend on Eσ^S^ dependent gene expression for survival (28).

## Material and Methods

### (p)ppGpp measurements

A culture of *E. coli* MG1655 *rpoC*-FLAG was setup from an overnight culture in 5 ml potassium morpholinopropane sulfonate (MOPS) minimal media with a starting OD_600_ of 0.05 at 37°C. At OD_600_ 0.1, 20 μCi/ml [^32^P] H_3_PO_4_ was added as phosphate source and the culture was left to grow to an OD_600_ of 0.45. The culture was then infected with T7 wild-type (ratio of 10:1 – T7:*E. coli*) in the presence of 1 mM CaCl_2_. To detect (p)ppGpp production, 500 μl of the cultures prior to infection (time 0) and at 10 min after infection were added to 100 μl of ice cold 2 M formic acid and incubated on ice for 30 min. The samples were then centrifuged for 5 min at 17,000x*g* and 10 μl of supernatant were spotted on thin layer chromatography (TLC) polyethylenimine (PEI) cellulose F (dimensions 20 cm x 20 cm [Merck Millipore]). The spot was left to migrate to the top of the sheet in a TLC tank in presence of 1.5 M KH_2_PO_4_ pH 3.6, dried before exposing overnight onto phosphor screen and viewed using Phosphoimager. For a positive control, 100 μg/ml of serine hyroxamate (SHX) was added to MG1655 *rpoC*-FLAG at OD_600_ of 0.45 and sample taken after 10 min and processed as above.

### Western blotting

*E. coli* MG1655 *rpoC*-FLAG strain was grown in LB at 30°C to an OD_600_ of ∼0.45. The culture was then infected with T7 wild-type (ratio of 0.1:1 – T7:*E. coli*) in the presence of 1 mM CaCl_2_. To detect σ^S^ production, 20 ml of the culture prior to infection (time 0) and at 10 min intervals after infection were taken until complete lysis was obtained. Experiments with the MG1655 Δ*relA*/Δ*spoT* strain ((29); kindly provided by Kenn Gerdes) were conducted exactly as described above but samples were taken at 0 and 40 min after T7 infection. Cultures were centrifuged, cell pellets re-suspended in 500 μl of 20 mM Na_2_PO_4_, 50 mM NaCl and 5% glycerol and sonicated. The cleared cell lysate was then loaded on 4-20% SDS-PAGE and ran at 200 V for 30 min. The SDS-PAGE gel was transferred onto polyvinylidene difluoride (PVDF) membrane (0.2 μm) using Trans-Blot^®^ Turbo™ Transfer System [Bio-Rad] device and processed according to standard molecular biology protocols. The primary antibodies were used at the following titres: anti-*E. coli* RNAP σ^S^ antibody at 1:500 [1RS1 – Biolegend], anti-*E. coli* RNAP α-subunit antibody at 1:1000 [4AR2 – Biolegend]. The secondary antibody Rabbit Anti-Mouse IgG H&L (HRP) was used at 1:2500 [ab97046––Abcam]. Bands were detected using an Amersham ECL Western Blotting Detection Reagent [GE Healthcare Life Sciences] and analysed on a Chemidoc using the Image Lab Software.

### Protein expression and purification

FLAG-tagged *E. coli* σ^70^ and σ^S^ were PCR amplified from *E. coli* genome and cloned into the pT7-FLAG^TM^-1 vector [Sigma-Aldrich]. Recombinant vectors pT7-FLAG::*rpoD* and pT7-FLAG::*rpoS* were confirmed by DNA sequencing. For the biochemical experiments, recombinant FLAG-tagged *E. coli* σ^70^ and σ^S^ were made by FLAG affinity purification from *E. coli* strain BL21 (DE3). Briefly, the culture of BL21 (DE3) cells containing pT7-FLAG::*rpoD* was grown at 37°C to an OD_600_ of □0.4, cold shocked on ice for 15 min before protein expression was induced with 0.1 mM IPTG. The cells were left to grow at 16°C overnight before harvesting. For FLAG-tagged *E. coli* σ^S^ expression, BL21 (DE3) containing pT7-FLAG::*rpoS* were grown at 37°C to an OD_600_ of L0.4 and protein expression was induced with 0.1 mM IPTG. The cells were left to continue growing at 37°C for 3 h before harvesting. The cell pellets for both FLAG-tagged *E. coli* σ^70^ and σ^S^ were re-suspended in binding buffer (50 mM Tris-HCl, 150 mM NaCl, pH 7.4) containing cocktail of protease inhibitors and lysed by sonication. The cleared cell lysate was loaded to a column containing anti-FLAG M2 affinity gel [Sigma-Aldrich] and the purified proteins were obtained by adding elution buffer (100 µg/ml 3XFLAG^®^ peptide [Sigma-Aldrich] in binding buffer) for 30 min at 4°C. The purified proteins were dialyzed into storage buffer (10 mM Tris-HCl pH 8.0, 50 mM NaCl, 50% glycerol, 0.1 mM EDTA and 1 mM DTT) and stored in aliquots at −80°C. The FLAG-tagged ^S^ΔR4 (amino acid residues 1-262) was made by introducing a stop codon into pT7-FLAG::*rpoS* by site-directed mutagenesis and its expression and purification was done as described for the full-length protein. The 6xHis-σ^S^R4 (amino acid residues 245-330) was amplified from *E. coli* genomic DNA by Gibson assembly and ligated into the pET-46 Ek/LIC vector [Merck Millipore] and expressed in *E. coli* strain BL21 (DE3) for the pull-down experiments and structural studies. The cells were grown in either LB (for pull-down experiments) or M9 Minimal medium labelled with ^15^N and ^13^C (for the structural studies) and induced with 0.5 mM IPTG when the OD_600_ reached 0.6 and incubated overnight at 18°C before harvesting by centrifugation. The cells were lysed by sonication in 50 mM NaH_2_PO_4_, 300 mM NaCl, 10 mM imidazole pH 8 and purified using Ni-NTA beads [Qiagen]. The eluate was then dialyzed against 50 mM NaH_2_PO_4_ and 350 mM NaCl, at pH 6 and subsequently concentrated down for NMR experiments whilst for pull-down experiments, the 6xHis-σ^S^R4 protein was kept in 50 mM NaH_2_PO_4_, 300 mM NaCl, and 10 mM imidazole pH 8. The 6xHis-σ^S^ was amplified from *E. coli* genomic DNA, cloned into pET-46 Ek/LIC vector [Merck Millipore] by Gibson assembly, expressed and purified as described for 6xHis-σ^S^R4. The cells were lysed by sonication under denaturing condition containing 8M urea, purified with Ni-NTA beads under denaturing conditions and the denatured 6xHis-σ^s^ was refolded in 50 mM NaH_2_PO_4_, 300 mM NaCl, and 10 mM imidazole pH 8. The 6xHis-Gp5.7 was amplified from pBAD18::*gp5.7* (8) and cloned into pET-46 plasmid. The Histidine tag on Gp5.7 was deleted to express tag-free Gp5.7 using Q5 site directed mutagenesis kit [NEB]. The protein was expressed under the same condition as 6xHis-σ^S^R4. Recombinant 6xHis-Gp5.7 and 6xHis-Gp2 expression and purification were done exactly as previously described (8, 10). Sequences of all oligonucleotides used in the construction of the expression vectors are available upon request.

### *In vitro* transcription assays

These were conducted exactly as previously described (8) in 10 mM Tris pH 7.9, 40 mM KCl, 10 mM MgCl_2_ using *E. coli* core RNAP from NEB and FLAG-tagged versions of σ^70^ and σ^S^ were purified exactly as described above. Reactions in Fig. 3C and 3D were conducted in 100 mM K-glutamate, 40 mM HEPES pH 8, 10 mM MgCl_2_ and 100 μg/ml BSA. In all reactions Gp5.7 or Gp2 was pre-incubated with Eσ^S^ or Eσ^70^ at the indicated concentrations before adding promoter DNA to the reaction. However, in the reactions shown in Fig. 3D, Gp5.7 was added to the pre-formed RP_O_ (i.e. following pre-incubation of Eσ^S^ and the promoter DNA). Sequences of all oligonucleotides used to generate promoter probes are available in (8) or upon request.

### Pull-down assays

For the pull-down assays shown in Fig. 2A and 2B, Ni-NTA beads [Qiagen] were used. Approximately 0.02 mg of recombinant 6xHis-σ^S^ or 6xHis-σ^S^R4 in binding buffer (50 mM NaH_2_PO_4_, 300 mM NaCl, and 10 mM imidazole pH 8) was added to beads and incubated at 4°C for 30 min. *E. coli* whole-cell lysate containing overexpressed untagged Gp5.7 was added to resin containing sigma and incubated for 1 h at 4°C. The beads were washed three times in 1 ml wash buffer (50 mM NaH_2_PO_4_, 300 mM NaCl, and 20 mM imidazole pH 8) for 10 min to remove any non-specific protein–protein interaction. To elute samples from beads, elution buffer containing 250 mM imidazole was added. For FLAG-tag protein pull-down assay (Fig. 2C), 0.02 mg FLAG-σ^S^ΔR4 was incubated with anti-FLAG M2 affinity gel [Sigma-Aldrich] in 50 mM Tris HCl, pH 7.4, 150 mM NaCl, 1 mM EDTA at 4°C for 2 h. *E. coli* whole-cell lysate containing overexpressed untagged Gp5.7 was added to resin containing sigma and incubated for 2 h at 4°C. The beads were washed three times in 50 mM Tris HCl, pH 7.4, 150 mM NaCl, 1 mM EDTA and 1% Triton X-100 for 10 min to remove any non-specific protein–protein interaction. To elute samples from beads, elution buffer containing 100 µg/ml 3XFLAG^®^ peptide [Sigma-Aldrich] was added. Ten microliters of samples together with Laemmli 2x concentrate SDS Sample Buffer was loaded on a 10–15% SDS-PAGE alongside Protein standard Marker and stained with Coomassie Brilliant Blue.

### NMR structure determination

NMR spectra were collected at 310K on Bruker DRX600 and DRX800 spectrometers equipped with cryo-probes. Spectral assignments were completed using our in-house, semi-automated assignment algorithms and standard triple-resonance assignment methodology (30). H_α_ and H_β_ assignments were obtained using HBHA (CBCACO)NH and the full side-chain assignments were extended using HCCH-total correlation (TOCSY) spectroscopy and (H)CC(CO)NH TOCSY. Three-dimensional ^1^H-^15^N/^13^C NOESY-HSQC (mixing time 100 ms at 800 MHz) experiments provided the distance restraints used in the final structure calculation (31). The ARIA protocol was used for completion of the NOE assignment and structure calculation. The frequency window tolerance for assigning NOEs was ±0.025 ppm and ±0.03 ppm for direct and indirect proton dimensions and ±0.6 ppm for both nitrogen and carbon dimensions. The ARIA parameters p, Tv and Nv were set to default values. 108 dihedral angle restraints derived from TALOS+ were also implemented. The 10 lowest energy structures had no NOE violations greater than 0.5 Å and dihedral angle violations greater than 5°. The structural statistics are shown in Supplementary Table 1.

### NMR titration

Unlabelled Gp5.7 was added to ^15^N labelled 6xHis-σ^S^R4 according to stoichiometric ratio to perform NMR titration. Maximum five-fold Gp5.7 was added to 6xHisσ^S^R4 in order to broad out the entire spectra.

### Electrophoretic mobility shift assays

These were conducted exactly as previously described to distinguish between initial promoter complex and RP_O_ formation (10). Briefly, 75 nM of *E. coli* core RNAP (NEB) was incubated with 300 nM σ^S^ or σ^S^ R4 either on ice (to monitor initial promoter complex formation) or at 37°C (to monitor RP_O_ formation) for 5 min in 100 mM K-glutamate, 40 mM HEPES pH 8, 10 mM MgCl_2_ and 100 μg/ml BSA. Twenty nanomolar of ^32^P-labelled T7 A1p was added and incubated for 5 min. Since initial T7 A1 promoter complexes (formed at temperatures < 4 °C) are sensitive to heparin and, conversely, RP_O_ (formed at 37°C) are resistant to heparin (32), the reactions were challenged with 100 μg/ml heparin before separating the RNAP bound and free promoter DNA by native gel electrophoresis on a 4.5% (w/v) native polyacrylamide gel run at 100 V for 100 min at 4°C (to monitor initial promoter complex formation) or for 60 min at room temperature (to monitor RP_O_ formation). The dried gel was then analyzed by autoradiography.

### Plaque-enlargement assay

T7 phage plaques were formed as described in (8). To obtain images of plaques, *E. coli* MG1655 cultures and MG1655 Δ*relA*/Δ*spoT* were grown to an OD_600_ of 0.45 in LB at 30°C and 300 μl aliquots of the culture were taken out and either T7 wild-type or T7 Δ*gp5.7* lysate (sufficient to produce ~ 10 plaques) were added together with 1 mM CaCl_2_ and incubated at 37°C for 10 min to allow the phage to adsorb to the bacteria. Three milliliters of 0.7% (w/v) top agar was added to each sample and plated onto plates containing exactly 20 ml of 1.5% (w/v) LB agar. The plates were then put in a Epson perfection V370 photo scanner [Model J232D] inside a 30°C incubator and images of the plates were taken every two hours over a 72 h period for analysis. For complementation experiments, *E. coli* MG1655 cells containing either pBAD18::empty or pBAD18::*gp5.7* or pBAD18::*gp5.7-L42A* (8) were used and the plaque-enlargement assay was carried out as above on plates containing 100 μg/ml ampicillin and 0.04% (w/v) L-arabinose to induce *gp5.7* expression. The MG1655 Δ*rpoS* strain was obtained by phage transduction from Δ*rpoS* mutant in Keio libray (33).

## ACKNOWLEDGMENTS

We would like to thank Daniel Brown for help with the (p)ppGpp measurements. This work was supported by Wellcome Trust awards to SM (Investigator Award 100280 and multiuser equipment grant 104833) and SW (Investigator Award 100958).

**Author contributions:**
A.T-S., B.L., S.W. and S.M. designed research; A.T-S., B.L., L.B., and D.B. preformed research; S.W., S.M., A.T-S., B.L., and U.Q. analyzed data and S.W., S.M., A.T-S and B.L. wrote the paper.

## References

1. De Smet J, Hendrix H, Blasdel BG, Danis-Wlodarczyk K, & Lavigne R (2017) Pseudomonas predators: understanding and exploiting phage-host interactions. Nat Rev Microbiol 15(9):517–530.

2. Nechaev S & Severinov K (2008) The elusive object of desire--interactions of bacteriophages and their hosts. Curr Opin Microbiol 11(2):186–193.

3. Nechaev S & Severinov K (2003) Bacteriophage-induced modifications of host RNA polymerase. Annu Rev Microbiol 57:301–322.

4. Kemp P, Gupta M, & Molineux IJ (2004) Bacteriophage T7 DNA ejection into cells is initiated by an enzyme-like mechanism. Mol Microbiol 53(4):1251–1265.

5. Savalia D, Robins W, Nechaev S, Molineux I, & Severinov K (2010) The role of the T7 Gp2 inhibitor of host RNA polymerase in phage development. J Mol Biol 402(1):118–126.

6. Severinova E & Severinov K (2006) Localization of the Escherichia coli RNA polymerase beta’ subunit residue phosphorylated by bacteriophage T7 kinase Gp0.7. J Bacteriol 188(10):3470–3476.

7. James E, et al. (2012) Structural and mechanistic basis for the inhibition of Escherichia coli RNA polymerase by T7 Gp2. Mol Cell 47(5):755–766.

8. Tabib-Salazar A, et al. (2017) Full shut-off of Escherichia coli RNA-polymerase by T7 phage requires a small phage-encoded DNA-binding protein. Nucleic Acids Res 45(13):7697–7707.

9. Friesen JD & Fiil N (1973) Accumulation of guanosine tetraphosphate in T7 bacteriophage-infected Escherichia coli. J Bacteriol 113(2):697–703.

10. Camara B, et al. (2010) T7 phage protein Gp2 inhibits the Escherichia coli RNA polymerase by antagonizing stable DNA strand separation near the transcription start site. Proc Natl Acad Sci U S A 107(5):2247–2252.

11. Liu B, Zuo Y, & Steitz TA (2016) Structures of E. coli sigmaS-transcription initiation complexes provide new insights into polymerase mechanism. Proc Natl Acad Sci U S A 113(15):4051–4056.

12. Chen H, Tang H, & Ebright RH (2003) Functional interaction between RNA polymerase alpha subunit C-terminal domain and sigma70 in UP-element- and activator-dependent transcription. Mol Cell 11(6):1621–1633.

13. Ross W, Schneider DA, Paul BJ, Mertens A, & Gourse RL (2003) An intersubunit contact stimulating transcription initiation by E coli RNA polymerase: interaction of the alpha C-terminal domain and sigma region 4. Genes Dev 17(10):1293–1307.

14. Minakhin L & Severinov K (2003) On the role of the Escherichia coli RNA polymerase sigma 70 region 4.2 and alpha-subunit C-terminal domains in promoter complex formation on the extended −10 galP1 promoter. J Biol Chem 278(32):29710–29718.

15. Sclavi B, et al. (2005) Real-time characterization of intermediates in the pathway to open complex formation by Escherichia coli RNA polymerase at the T7A1 promoter. Proc Natl Acad Sci U S A 102(13):4706–4711.

16. Zuo Y & Steitz TA (2015) Crystal structures of the E. coli transcription initiation complexes with a complete bubble. Mol Cell 58(3):534–540.

17. Dominguez C, Boelens R, & Bonvin AM (2003) HADDOCK: a protein-protein docking approach based on biochemical or biophysical information. J Am Chem Soc 125(7):1731–1737.

18. Battesti A, Majdalani N, & Gottesman S (2011) The RpoS-mediated general stress response in Escherichia coli. Annu Rev Microbiol 65:189–213.

19. Nowicki D, Kobiela W, Wegrzyn A, Wegrzyn G, & Szalewska-Palasz A (2013) ppGpp-dependent negative control of DNA replication of Shiga toxin-converting bacteriophages in Escherichia coli. J Bacteriol 195(22):5007–5015.

20. Yin J (1993) Evolution of bacteriophage T7 in a growing plaque. J Bacteriol 175(5):1272–1277.

21. Farewell A, Kvint K, & Nystrom T (1998) Negative regulation by RpoS: a case of sigma factor competition. Mol Microbiol 29(4):1039–1051.

22. Bae B, et al. (2013) Phage T7 Gp2 inhibition of Escherichia coli RNA polymerase involves misappropriation of sigma70 domain 1.1. Proc Natl Acad Sci U S A 110(49):19772–19777.

23. Hinton DM (2010) Transcriptional control in the prereplicative phase of T4 development. Virol J 7:289.

24. North SH, Kirtland SE, & Nakai H (2007) Translation factor IF2 at the interface of transposition and replication by the PriA-PriC pathway. Mol Microbiol 66(6):1566–1578.

25. Slominska M, Neubauer P, & Wegrzyn G (1999) Regulation of bacteriophage lambda development by guanosine 5’-diphosphate-3’-diphosphate. Virology 262(2):431–441.

26. Hauryliuk V, Atkinson GC, Murakami KS, Tenson T, & Gerdes K (2015) Recent functional insights into the role of (p)ppGpp in bacterial physiology. Nat Rev Microbiol 13(5):298–309.

27. Robertson ES, Aggison LA, & Nicholson AW (1994) Phosphorylation of elongation factor G and ribosomal protein S6 in bacteriophage T7-infected Escherichia coli. Mol Microbiol 11(6):1045–1057.

28. Dong T & Schellhorn HE (2010) Role of RpoS in virulence of pathogens. Infect Immun 78(3):887–897.

29. Magnusson LU, Gummesson B, Joksimovic P, Farewell A, & Nystrom T (2007) Identical, independent, and opposing roles of ppGpp and DksA in Escherichia coli. J Bacteriol 189(14):5193–5202.

30. Marchant J, Sawmynaden K, Saouros S, Simpson P, & Matthews S (2008) Complete resonance assignment of the first and second apple domains of MIC4 from Toxoplasma gondii, using a new NMRView-based assignment aid. Biomol NMR Assign 2(2):119–121.

31. Pardi A (1995) Multidimensional heteronuclear NMR experiments for structure determination of isotopically labeled RNA. Methods Enzymol 261:350–380.

32. Schickor P, Metzger W, Werel W, Lederer H, & Heumann H (1990) Topography of intermediates in transcription initiation of E.coli. EMBO J 9(7):2215–2220.

33. Baba T, et al. (2006) Construction of Escherichia coli K-12 in-frame, single-gene knockout mutants: the Keio collection. Mol Syst Biol 2:20060008.

